# Bub1 is not required for the checkpoint response to unattached kinetochores in diploid human cells

**DOI:** 10.1101/278820

**Authors:** Cerys E. Currie, Mar Mora-Santos, Chris Smith, Andrew D. McAinsh, Jonathan B.A. Millar

**Affiliations:** Centre for Mechanochemical Cell Biology & Division of Biomedical Sciences, Warwick Medical School, University of Warwick, Gibbet Hill, Coventry, CV4 7AL, UK; Present address: Metabolic Research Laboratories and MRC Metabolic Diseases Unit, Institute of Metabolic Science, Addenbrooke’s Hospital, Hills Road, Cambridge CB2 0QQ, UK

## Abstract

Error-free chromosome segregation during mitosis depends on a functional spindle assembly checkpoint (SAC). The SAC is a multi-component signaling system that is recruited to incorrectly attached kinetochores to catalyze the formation of a soluble inhibitor, known as the mitotic checkpoint complex (MCC), which binds and inhibits the anaphase promoting complex [1]. We have previously proposed that two separable pathways, composed of KNL1-Bub3-Bub1 (KBB) and Rod-Zwilch-Zw10 (RZZ), recruit Mad1-Mad2 complexes to human kinetochores to activate the SAC [2]. We refer to this as the dual pathway model. Although Bub1 is absolutely required for MCC formation in yeast (which lack RZZ), there is conflicting evidence as to whether this is also the case in human cells based on siRNA studies [2–5]. Here we report, using genome editing, that Bub1 is not strictly required for the SAC response to unattached kinetochores in human diploid hTERT-RPE1 cells, consistent with the dual pathway model.

## Results and Discussion

We used CRISPR/Cas9 genome editing to disrupt both alleles of *BUB1* in human diploid hTERT-RPE1 cells using small guide (sg)RNAs targeting exon 2. Genome sequencing revealed a frame shift in both *BUB1* alleles that allows expression of only the first 23 amino acids of Bub1 (Figure S1). This was confirmed by immunoblotting and quantitative immunofluorescence using multiple Bub1 antibodies (Figure 1A,B). Importantly, Rod and Zwilch (subunits of RZZ complex), as well as KNL1, bound kinetochores to the same extent in parental and Bub1^-/-^ cells (Figure 1C and S2A). By contrast, BubR1 kinetochore-binding was abolished and binding of CENP-F severely compromised (Figure 1C, S2A), as reported previously [6]. Importantly, steady state levels of Mad2 at pro-metaphase kinetochores were lower in Bub1^-/-^ cells than in the parental control (Figure 1C, S2A), consistent with previous findings following siRNA mediated knockdown of KNL1 or Bub1 [2].

**Figure 1.**
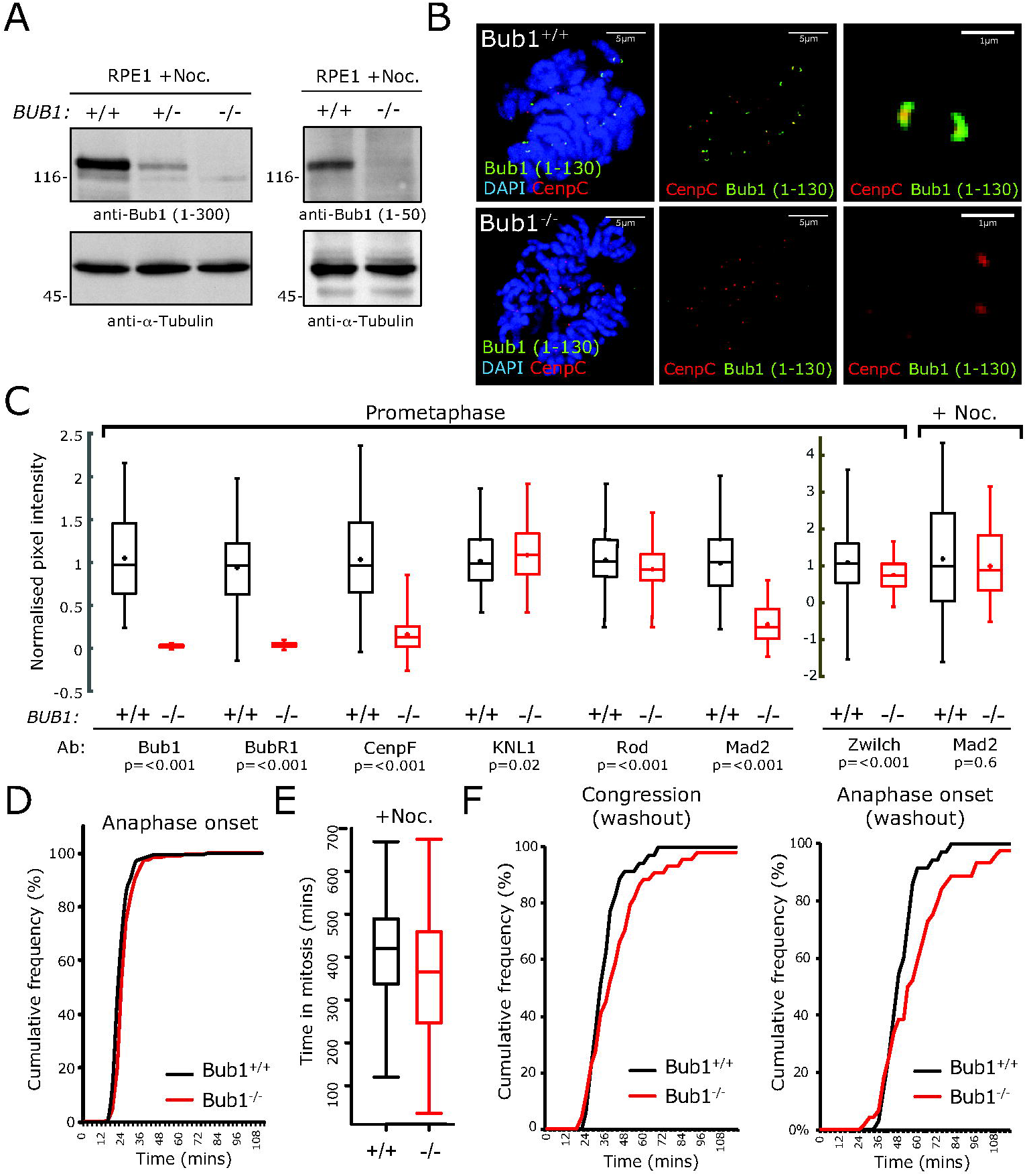
BUB1 is not absolutely required for SAC activity in diploid human cells. (A) lmmunob lots of whole-cell lysates from parental or Bub1^-/-^ cells treated with 3.3 µM nocodazole and probed with antibodies against Bub1 aa1-300 (left) and aa1-50 (right), a-Tubulin as a loading control.(B) Representative images of prometaphase parental or Bub1^-/-^ cells stained with DAPI, CenpC, and Bub1 (aa1-130) antibodies. Scale bar = 5 µm. (C) Quantification of kinetochore-bound Bub1, BubR1, CenpF, KNL1, Rod, Zwilch and Mad2 signals relative to Crest or CenpC intensity (n= >200 kinetochores; data normalised to respective Bub1^+/+^ median value; p-value from a two-sided Mann-Whitney U test). Data from >1 independent experiments. Representative images are shown in Figure S1. (D) Quantification of time spent in mitosis (from NEB to anaphase onset or mitotic exit from live-cell videos of parental (n=183) or Bub1-/- (n= 218) cells treated SiR-DNA (to visualise DNA), from >3 independent experiments. (E) Same experiment as (D) except cells treated with 100nM nocodazole (n=66 for Bub1^-/-^ and n=19 for parental). Data from >1 independent experiments. (F) Quantification of mitotic progression following nocodazole washout. Timing of washout to completion of chromosome congression (left), and washout to anaphase onset (right) in parental (n=53) or Bub1^-/-^ (n= 62) cells. Data from >4 independent experiments. Representative images are shown in Figure S1.

We next used live cell imaging of chromosomes labeled with a cell permeable dye (SiR-DNA) to assess the effect of Bub1 knockout on timing of both anaphase onset and chromosome congression (time until last chromosome aligns in metaphase plate) in individual cells. To our surprise, we found that the efficiency of congression was unaffected in Bub1^-/-^ cells compared to control cells, whilst the time from nuclear breakdown (NBD) to anaphase onset was extended by ∼3 min (Figure 1D). Since depletion of MCC by siRNA knockdown of Mad2 or BubR1 shortens the NBD-to-anaphase period [7], these results suggest that Bub1 is not required for MCC formation in an unperturbed mitosis. To directly test whether Bub1 is required for the SAC response to unattached kinetochores we filmed control and Bub1^-/-^ cells in the presence of 100nM nocodazole and measured the time from NBD to anaphase onset. We found that addition of nocodazole delays anaphase onset in Bub1^-/-^ cells to a similar extent as in control cells (Figure 1E). Moreover, we found that Mad2 binds unattached kinetochores (Figure 1C, far right panel) and interacts with BubR1, Cdc20 and APC/C almost as efficiently in checkpoint arrested Bub1^-/-^ cells as the parental control (Figure S3A). This shows that Bub1 is neither required for the formation of MCC or MCC-APC/C, nor the checkpoint response to unattached kinetochores. It also implies that Bub1 is not required for MCC formation from either the RZZ complex or the nuclear pore [8]. Furthermore, these data also indicate that recruitment of BubR1 to kinetochores is not necessary for its incorporation into MCC or for checkpoint signaling.

The apparent absence of a congression defect in Bub1^-/-^ cells is consistent with previous work in RPE1 cells with Bub1 kinase inhibitors [9], but is surprising given the proposed role of Bub1 in error correction through the Histone2A-Sgo1/PP2A-Aurora B pathway [10]. This may be due to the rapid bi-orientation and alignment of kinetochores in RPE1 cells [11]. We therefore released cells from a nocodazole arrest, generating multiple mal-orientated attachments requiring correction and bi-orientation, and measured the time to completion of congression and anaphase onset. Both these events were delayed in Bub1^-/-^ cells compared to the control (15 min delay in 80% cells congressing, and 9 min delay in 80% cells reaching anaphase), revealing the role for Bub1 in error correction (Figure 1F, S2B). Consistently, association of Sgo1 to the centromeric region between separated sister kinetochores is reduced by approximately 50% in Bub1^-/-^ cells (Figure S3B), similar to that observed previously using depletion of Bub1 by siRNA in HeLa cells [10]. Interestingly, the frequency of lagging chromosomes was unchanged in Bub1^-/-^ compared to parental cells (19% vs. 10%, p=0.184 Fisher’s exact test) suggesting that while the efficiency of error correction is reduced, Bub1^-/-^ cells are still able to successfully complete bi-orientation. Moreover, Bub1^-/-^ cells with uncongressed chromosomes delay in mitosis, consistent with our conclusion that the SAC can operate without Bub1. We assume that error correction is completed in Bub1^-/-^ cells by a pool of Aurora-B kinase associated with centromeric DNA via phosphorylation of histone H3 on threonine 3 (H3T3) by cohesion-bound Haspin kinase [12].

The data in this paper support the hypothesis that KBB and RZZ complexes provide two separate receptors for the Mad1-Mad2 complex at human kinetochores (the dual pathway model [2]). However, it is presently unclear why human cells have two receptors (KBB and RZZ), whereas yeast only has one (KBB). One possibility is that the KBB and RZZ pathways enable monitoring of different attachment states. Our previous work in human cells hints at a role for the KBB pathway in delaying anaphase onset when chromosomes are either not fully attached and/or not properly aligned at the metaphase plate [2]. The generation of Bub1^-/-^ cells allows such models to be rigorously tested. The key challenge will be to determine how the KBB and RZZ complexes coordinate checkpoint signaling and silencing. Importantly, these data are entirely consistent with a recent report showing that Bub1 is not essential for the spindle checkpoint response to unattached kinetochores in near-haploid human HAP1 cells, which are derived from a male patient with chronic myelogenous leukemia (CML) [13]. It is important to point out, however, that the contribution of Bub1 and the KBB pathway to SAC signaling and error correction may differ between cell types, developmental stages or stages of cellular transformation. Indeed, knockdown of KBB by siRNA has a more penetrant (congression) phenotype in transformed aneuploid HeLa cells than in RPE1 cells [2,7], and experiments in mouse models point to an essential role for Bub1 during development [14]. More particularly, recent data reveal an altered requirement for the KBB pathway following the epithelial-to-mesenchymal transition [15]. Future work will be needed to understand whether the roles of KBB and RZZ in SAC signaling and error correction are coupled to normal development and cancer progression.

## SUPPLEMENTAL INFORMATION

Supplemental Information includes experimental procedures and three supplemental figures.

## AUTHOR CONTRIBUTIONS

This project was initiated by A.D.M. and J.B.M. C.E.C. conducted all experiments with the exception of biochemistry in Figure 2A and Figure S3A, which was carried out by M.M.S. C.S. wrote the code for quantification of kinetochore signals. The manuscript was written by A.D.M., J.B.M., and C.E.C.

## ACKNOWLEDGEMENTS

We thank Muriel Erent for support in genome editing and V. Silió, John Meadows and Steve Royle for comments on the manuscript. This work was supported by MRC program grant MR/K001000/1 to A.D.M. and J.B.M. C.E.C. is funded by the BBSRC Midlands Integrative Biosciences Training Partnership (*MIBTP*) [grant BB/M01116X/1]. A.D.M and C.S. were also supported by a Wellcome Trust Senior Investigator Award (grant 106151/Z/14/Z) and a Royal Society Wolfson Research Merit Award (grant WM150020).

---

**Figure S1:**
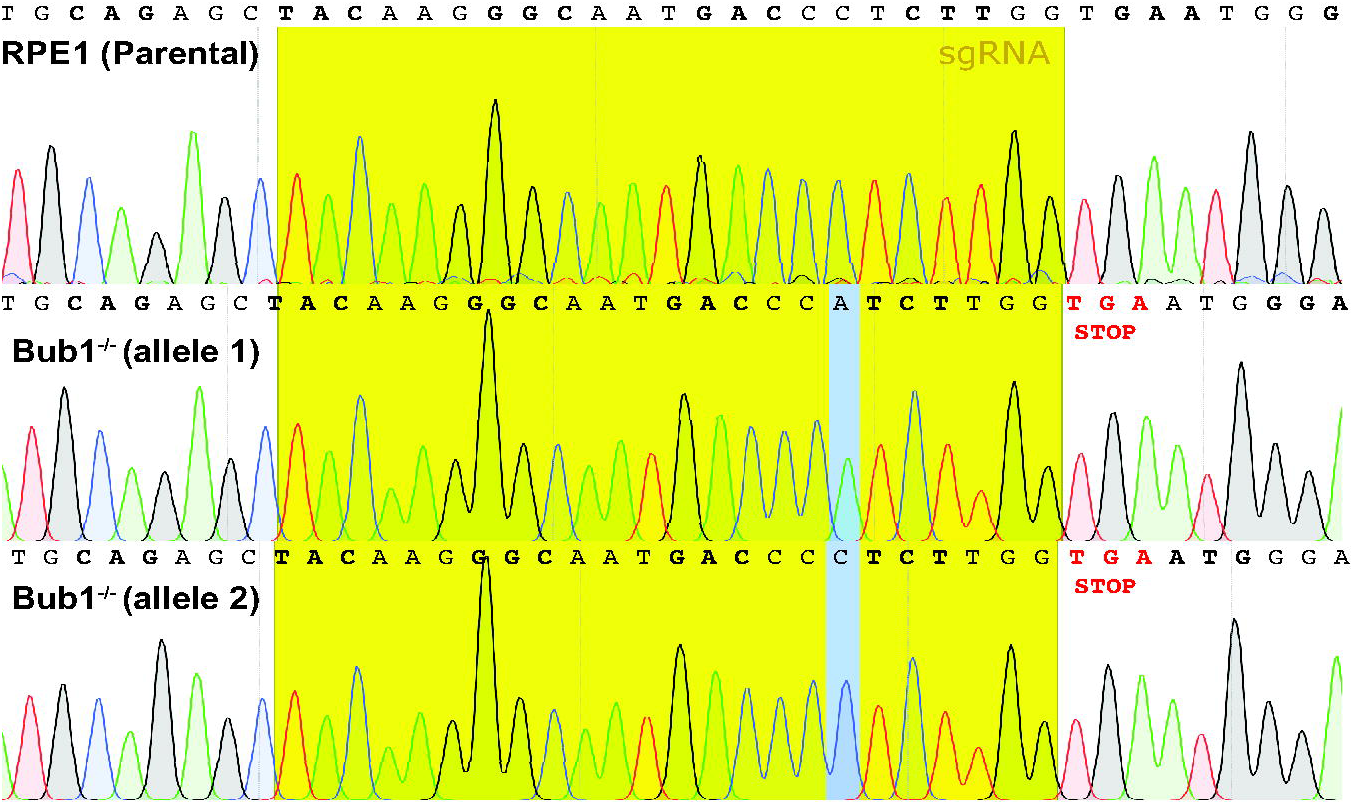
Genome sequencing of Bub1^-/-^ cells showing a frameshift mutation in both BUB1 alleles due to the addition of 1 base pair. The frameshift induces premature stop codons in both alleles.

**Figure S2:**
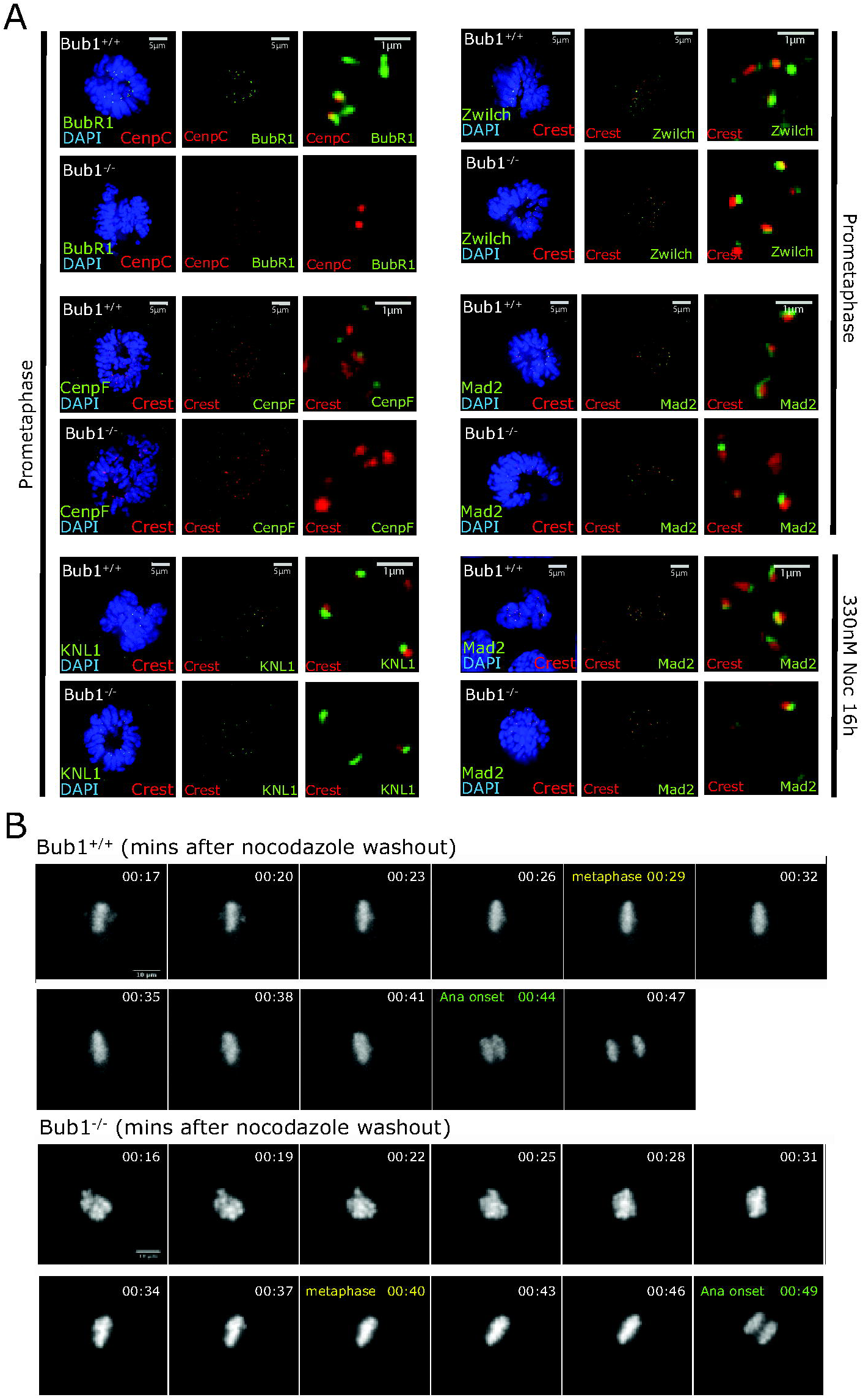
(A) Representative images of prometaphase or Nocodazole treated parental and Bub1^-/-^ cells stained with DAPI, Crest/CenpC, and BubR1, CenpF, KNL1, Rod, and Mad2 antibodies. For Quantification see Figure 1C. (B) Representative stills from live-cell videos of parental or Bub1^-/-^ cells treated SiR-DNA (to visualise DNA) following nocodazole washout. Time in h:min.

**Figure S3:**
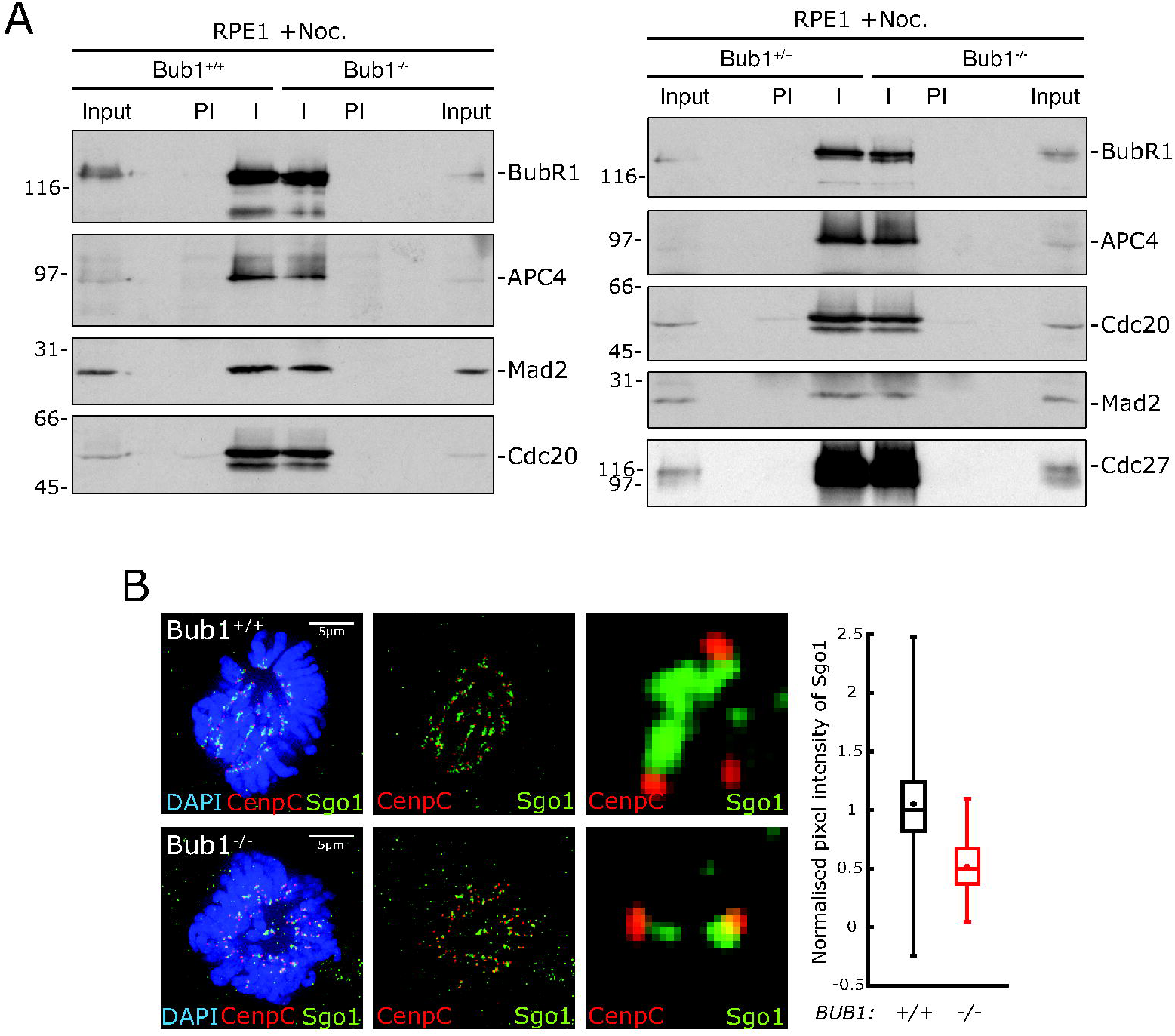
(A) Assembly of Mitotic checkpoint complex (MCC) in absence of Bub1. Parental or Bub1^-/-^ cells were treated with 3.3 μM Nocodazole for 24 hours. Mitotic cell extracts (Input) were im mu no precipitated with mouse anti-Cdc20 or anti-APC3 (Immune; I) or normal mouse serum (Pre-lmmune; PI) and complexes analysed by immunoblotting with antibodies as indicated. (B) Sgo1 localisation to centromeric NDA is substantially reduced in the absence of Bub1. Representative images of prometaphase parental and Bub1^-/-^cells stained with DAPI, CenpC, and Sgo1 antibodies. Quantification of Sgo1 levels at kinetochores, normalised to CenpC intensity.

